# Linking cross-species trajectories of cerebrovascular remodeling in aging and Alzheimer’s disease to brain vessel transcriptome

**DOI:** 10.64898/2026.03.06.710077

**Authors:** Noah Schweitzer, Yucheng Shen, Yifan Zhao, Christopher Cover, Shahnur Alam, Rebecca Deek, Jiatai Li, George Stetten, Howard Aizenstein, Minjie Wu, Rada Koldamova, Alberto Vazquez, Nicholas F. Fitz, Bistra Iordanova

## Abstract

Cerebrovascular remodeling driven by subtle molecular changes starts early in the asymptomatic stage of Alzheimer’s disease (AD). Despite progress in human vascular imaging and *postmortem* tissue analysis, there is limited data on the early features of small vessel reorganization, particularly in the context of cell-specific molecular drivers. This is largely because of the invasive nature of the tools for direct cellular observation and analysis. Since early detection is key, histopathology falls short with end-point data from people that died in late stages of the disease. This is a critical knowledge gap, because the early vascular processes are thought to be strongly correlated with health outcomes, tipping the scales from mild cognitive impairment to AD. To meet these translational challenges, we performed near life-span *in vivo* two-photon imaging and MRI of the cerebrovascular tree in a mouse model of amyloidosis. We identified precisely when subtle abnormalities in vessel tortuosity and red blood cell velocity first emerge in the context of differential amyloid accumulation in vessels walls and tissues. We then isolated the brain vessels for transcriptional analysis at this flagship timepoint and performed cross-species analysis linking changes in vascular cells to genes and pathways common to both mice and humans. Importantly, using 7T MRI of aging humans, we directly associated vascular remodeling trajectories of mice and humans and identified a remarkably analogous tortuosity course in the smallest brain vessels. Our integrated framework across scales and species advances neuroimaging biomarker understanding and uncovers early mechanistic routs of dysfunctional angiogenesis and actin-mediated contractility.

## INTRODUCTION

Alzheimer’s disease (AD) is defined by neuropathological measures of amyloid β (Aβ) plaques accumulation, followed by tau tangles, neurodegeneration and ultimately, cognitive decline and dementia (*1*). Breakthroughs of non-invasive amyloid imaging improved *perimortem* diagnosis, but they also revealed that the AD continuum spans decades beginning with prodromal brain changes that are unnoticeable by the affected person (*2*). It is still not clear how long each person spends in each part of the AD continuum, however, it is certain that some individuals maintain cognitive function longer, despite high amyloid load, making amyloid imaging a relatively weak predictor of conversion to AD (*3*). Several converging lines of evidence implicate dysfunctional brain vasculature as a chief contributor to the early AD pathophysiology (*4, 5*). Studies in humans and rodents report decreased cerebral blood flow, chronic hypoxia, and compromised blood-brain barrier coinciding with early Aβ deposition in asymptomatic subjects (*6–8*). It is not clear whether the amyloid in the walls of brain vessels, termed cerebral amyloid angiopathy (CAA), plays a distinct role, because there are no clinical non-invasive tests specific for CAA. In rodents, techniques like multi-photon microscopy offer high resolution vessel and CAA imaging but remain limited to the cortex. Conversely, time-of-flight (TOF) MRI can probe whole brain vascular trees, but lacks the resolution to image the capillary bed (*9*). Furthermore, accurate vessel segmentation remains a key analytical bottleneck (*10*). Finding vascular biomarkers in the context of CAA and plaques progression is becoming even more important as pressure increases to develop early AD treatments targeted appropriately.

Decreased vascular density in both AD humans (*11, 12*) and mouse models (*13, 14*) is well-documented, however transcriptomic analyses display angiogenic and pro-inflammatory signaling which is discordant with low vascular density, but it could be linked to the tortuous small vessels seen in histology of AD brains (*12, 15*). *In vitro* evidence indicates that vessel tortuosity exerts sheer stress on the endothelium and can trigger inflammation and aberrant angiogenesis (*16*). The *in vivo* temporal order these events in the early AD stages is still unclear, because human vascular transcriptome is acquired *postmortem* and thus limited to static measures from individuals with advanced AD. These methodological constraints limit information on cellular level during early AD stages and fail to capture events occurring at the critical period when vascular abnormalities emerge in asymptomatic subjects. In effort to fill these knowledge gaps and overcome the existing technological limitations, we carried a multimodal study in mice and humans to identify changes across the entire cerebrovascular network during normal aging and AD. We aimed to overcome the discontinuity between single-cell and system-level discoveries by integration of optical imaging and MR angiography with analysis of transcriptional vascular signatures across scales and species. We imaged longitudinally male and female mice with progressive amyloidosis to determine when cerebrovascular abnormalities first emerge and then we identified the corresponding transcriptional landscape. We isolated cerebral vessels for bulk RNA-seq, which yielded differentially expressed genes clustered around cellular pathways aligned with our neuroimaging observations. To translate the AD animal model findings, we first mapped the identified genes to a healthy mouse brain vascular atlas (*17*), and then to healthy and AD human brain vascular atlases (*18*). To validate the human relevance of cerebrovascular tortuosity as a biomarker, we imaged the arterial tree of cognitively normal older adults using ultra-high-field 7T MRI. We observed a striking similarity between mice and humans in the increases of age-related tortuosity of the small, but not the large arteries, supporting its potential as an early marker of vascular aging and vulnerability to AD. Our cross-species study identified transcriptomic signatures underlying vascular abnormalities observed through imaging at a critical time when vascular remodeling and blood flow dysfunction first emerge in AD. Such vulnerability signatures can serve as therapeutic targets if they can be altered, or as biomarkers if they are accessible. Our findings are also particularly valuable for capitalizing on existing human MRI measures during the early stages of AD.

## RESULTS

### Vascular morphology aging trajectories in mouse models of amyloidosis

To assess brain microvascular changes, we conducted longitudinal two-photon imaging of awake mice ages 3-18 months through a chronic cranial window with blood vessels labeled *in vivo*. We used 18 AD mice (B6.Cg-TgAPPswe, PSEN1dE9) and 18 control (B6.C3F1/J), 1:1 male and female. We trained a 5-layer residual U-net model to segment the vasculature (Fig. 1A), which was then skeletonized to quantify vessel tortuosity and junction density (Fig. 1B). We observed an early increase in microvascular tortuosity for AD mice that started before 6 months, peaked at 7-11 months (*p* < 0.001) and plateaued after 12 months (Fig. 1C). Control mice had no significant difference between <=6 months and 7-11 months, but then significantly increased after 12 months (*p* < 0.001). The mean tortuosity we computed was in line with previous reports (*14, 19*). Linear regression analysis revealed junction density decreased with age for AD mice (*p* = 0.0031), but no change in controls (control Age, *p* = 0.80; Age*Model, *p* = 0.045). No sex differences were observed in tortuocity.

**Figure 1.**
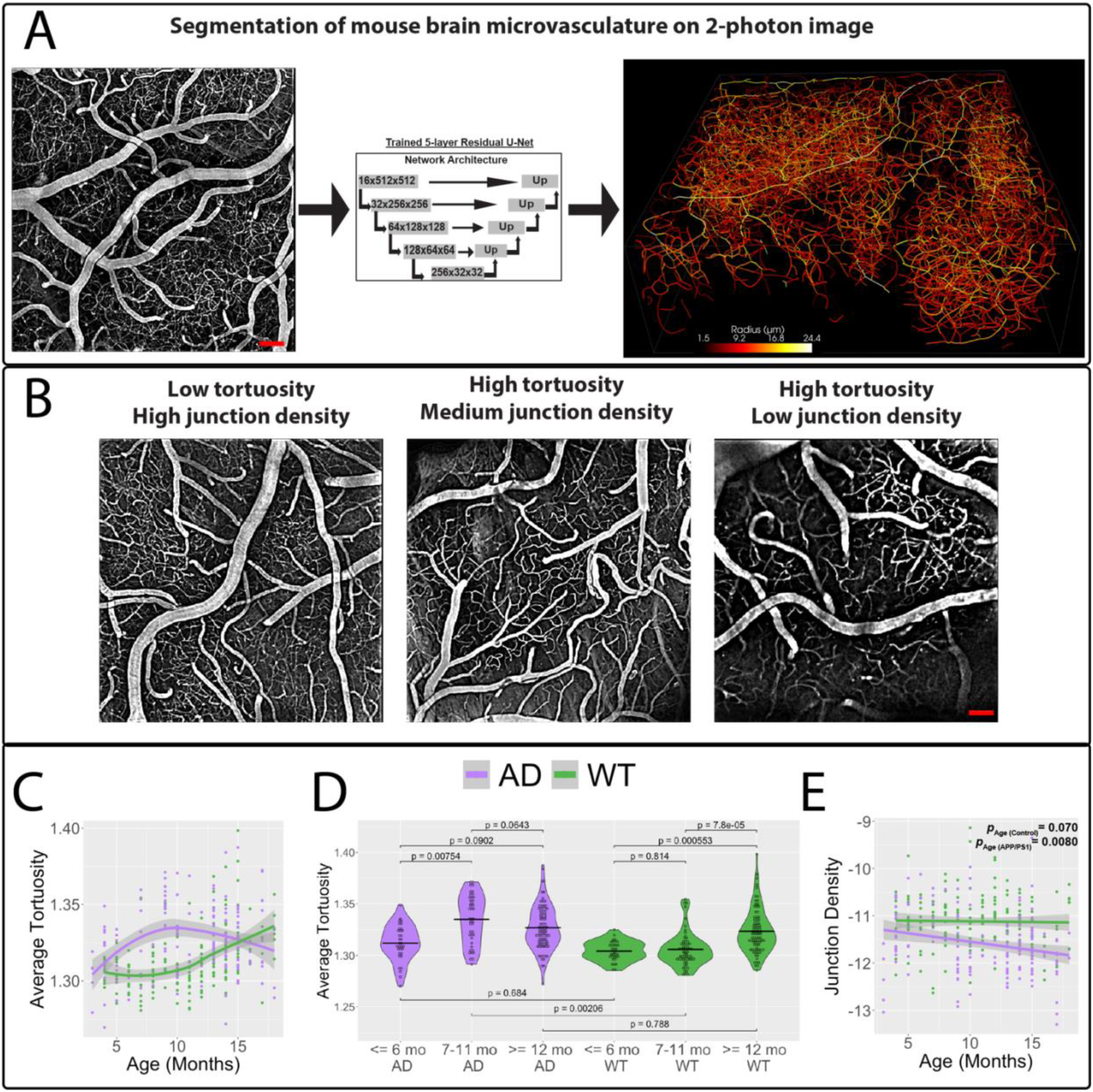
Near-lifespan longitudinal tracking of mouse brain microvasculature using 2-photon microscopy. A.) A 5-layer residual U-Net deep learning model was trained to segment the mouse brain microvasculature from 2-photon microscopy images (N = 9 per model and per sex, aged 3-18 months). Scale bar = 20 um. B.) Vessel tortuosity and junction density were quantified in each segmented vascular tree. Scale bar = 20 um. C.) LOESS-based trajectory of brain microvascular tortuosity showing that tortuosity increases early in life for AD mice while controls experience an increase much later. D.) In AD mice, tortuosity significantly increases around 7 months and plateaus after 11 months, whereas in controls, tortuosity begins to increase around 12 months. Analysis by ANOVA followed by Tukey’s multiple comparison test; bars represent mean ± SEM. *p < 0.05; **p < 0.01; ***p < 0.001. E.) Linear regression analysis reveals junction density in AD mouse brain vasculature decreases significantly compared to controls. Green and purple p-values represent the results of linear regression analyses examining the relationship between junction density and age for AD and WT mice, respectively. In panels (C–D), each data point corresponds to a 3D two-photon image volume acquired during a single session per mouse. N = 160 APP/PS1 (65 females and 95 males), 176 WT (51 females and 125 males). ∗∗∗∗p < 0.0001, ∗∗p < 0.01, ∗p < 0.05.

To assess whole-brain cerebrovascular changes, we imaged the large arterial tree in a separate cohort of AD (N = 47; 23 male, 24 female) and B6C3 control mice (N = 17; 8 male, 9 female) aged 2-25 months using TOF MRI. We trained a second 5-layer residual U-net model to segment the large arterial tree (Fig. 2A and B) which was registered to a B6C3 MRI atlas (*20, 21*) (Fig 2C). No significant changes in large vessel tortuosity were observed with aging in either AD (males p_age_ = 0.911, females p_age_=0.289) or controls (males p_age_ = 0.691, females p_age_=0.156; Fig. 2D). The only significant regional effect was an age-by-sex interaction in the subcortex, where tortuosity increased with age in AD males, but not in AD females or control mice (*p_age*sex_* = 0.0022; Fig. 2E). Junction density in the large arterial tree was significantly higher in males than females, controlling for model and age (AD *p_sex_* = 0.023; control *p_sex_* = 0.20; all *p_sex_* = 0.0092; Fig. 2F). The total vessel density was lower in AD mice compared to controls for the hypothalamus (*p* = 0.0037), but no other region in brain had significant model differences after multiple comparison correction. The total vessel density was higher in male mice compared to females, regardless of age and model, for the total brain (*p* = 0.0054) along with the cortex, subcortex, thalamus, olfactory bulb, hippocampus, caudate and amygdala (*p* < 0.05). Combined, these optical and MRI findings suggest that the microvasculature undergoes significant changes early in AD progression whereas the large arteries begin changing later in life. Importantly, in addition to age, we found that sex is an important factor for vascular density across the lifespan.

**Figure 2.**
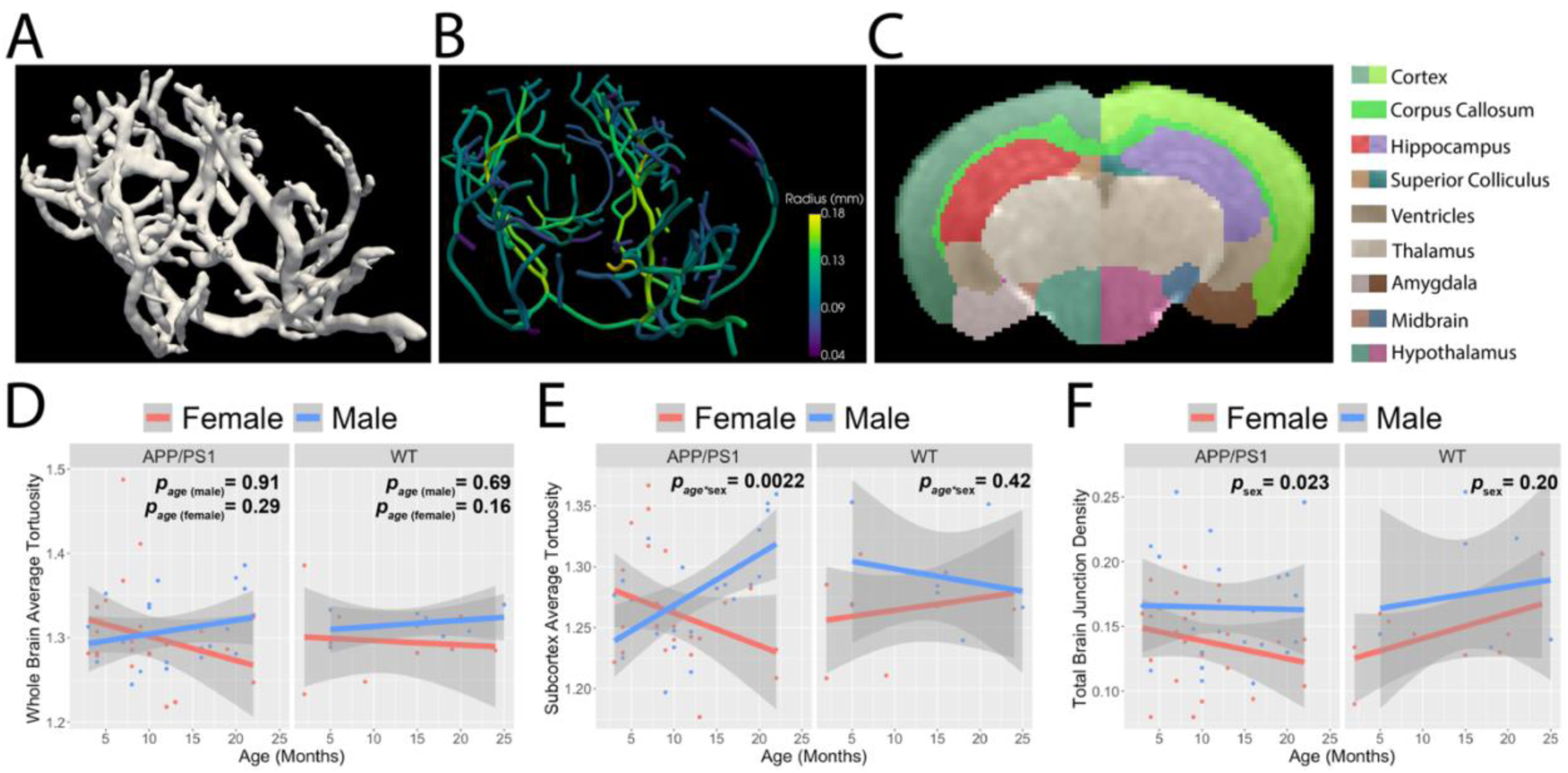
Sex and region-dependent differences in the vascular tree of aging mice. A.) A 5-layer residual U-Net deep learning model was trained to segment large arteries in the mouse brain from time-of-flight MRI (N = 10 per model and per sex, aged 2-25 months). B.) The segmented large arteries were then skeletonized for subsequent analysis. C.) Mouse brains were registered to an atlas (*20, 21*) to probe regional changes in large arteries. D.) Whole-brain large artery tortuosity does not significantly change with aging in either group AD (males p_age_ = 0.911, females p_age_=0.289) and WT (males p_age_ = 0.691, females p_age_=0.156 . E.) Subcortical large artery tortuosity increases only in AD male mice (multivariate linear regression analysis; AD mice interaction effect Age*Sex *p* = 0.0022). F.) Whole-brain large artery junction density is greater in males compared to females, irrespective of mouse model and age (multivariate linear regression analysis; AD *p_sex_* = 0.023;WT *p_sex_* = 0.20; All *p_sex_* = 0.0092). In panels (D–F), red and blue p-values indicate the results of linear regression analyses assessing the relationship between the TOF-derived measures on the y-axis and age for AD and WT mice, respectively. Each data point corresponds to a 3D time-of-flight MR image volume acquired during a single session per mouse. N = 47 APP/PS1 (24 females and 23 males), 17 WT (9 females and 8 males).

#### Resting capillary blood flow across healthy aging and AD

In addition to longitudinal two-photon imaging of the microvasculature morphology, we also quantified the capillary blood flow rate from 2,373 vessels in the same aging cohort. Capillary blood flow rate was derived from red blood cell (RBC) velocity and vessel diameter (Fig. 3A). RBC velocity was computed using Radon transform of line-scans (*22, 23*) (Fig. 3B) and the vessel diameter was measured using the full-width-half-maximum of the intensity profile of vessel cross-section (Fig. 3C). We observed a significant decrease during aging in the capillary blood flow rate for AD mice (*p* = 3.2E-05) and no change for controls (Age *p* = 0.71; interaction effect Age*Model *p* = 4.7E-8; Fig. 3D).

**Figure 3.**
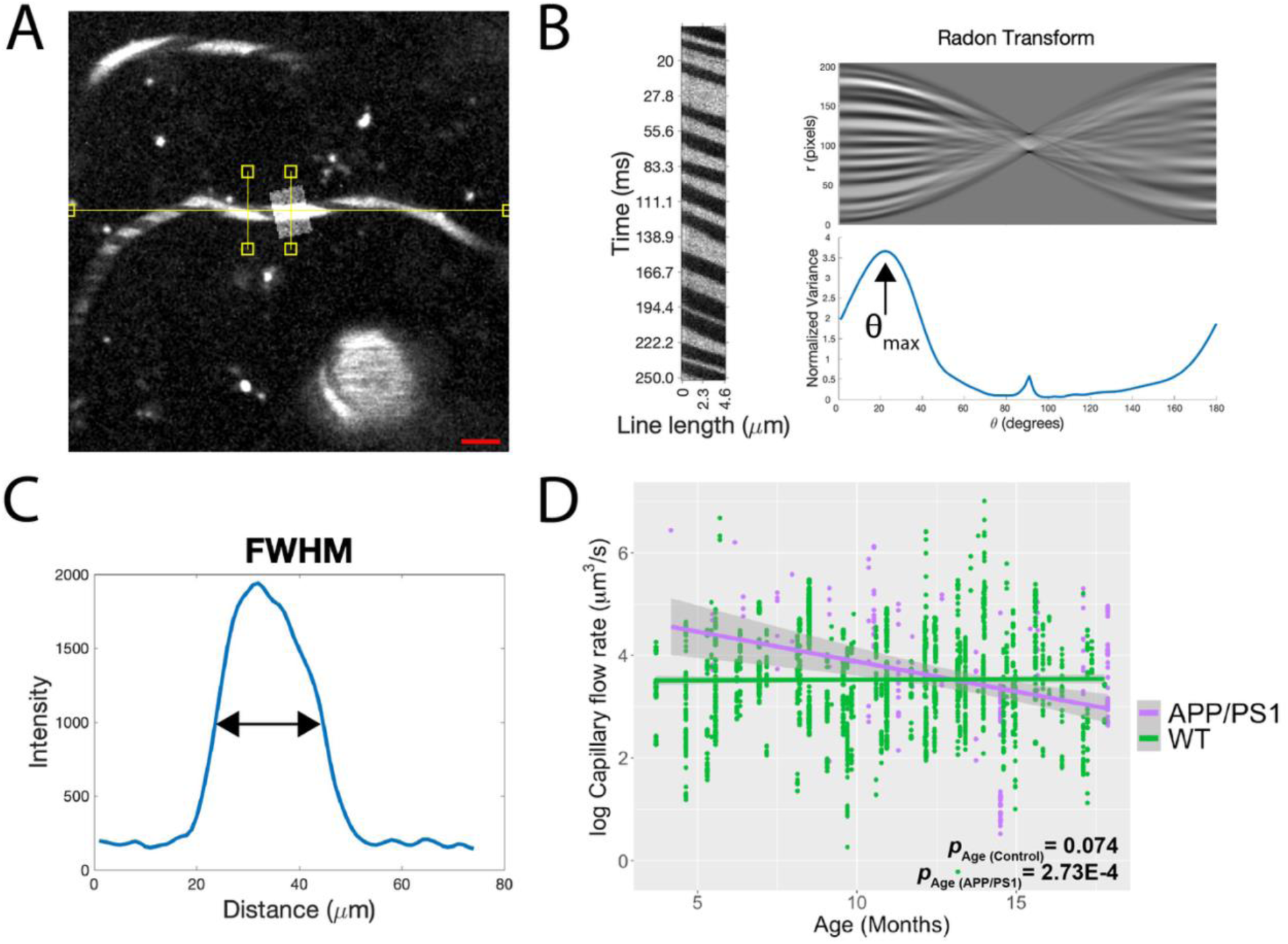
Near-lifespan tracking of capillary blood flow from measurement of 2,373 vessels displays a pronounced decrease with age in AD mice. A.) Capillary blood flow rate was calculated by measuring diameter (shaded rectangle) and red blood cell velocity (yellow line) in 2,373 capillaries across different ages in AD and control mice (N = 9 per model and per sex, aged 4-18 months). Scale bar = 20 um. B.) Red blood cell (RBC) velocity was measured using line-scan imaging. Each row in the corresponding line scan image (left) represents a single time point. The angle of the dark stripes indicates RBC speed. The Radon transform was applied to the line-scan image (top-right), and the angle that returns the largest variance was selected to compute RBC velocity (bottom-right). C.) Capillary diameter was calculated using the full-width half maximum of the vessel cross-section intensity. D.) Capillary flow rate significantly decreases with age in AD mice but not controls. Multivariate linear regression analysis; age for AD mice *p* = 3.2E-05 (purple p-value) ; age for control mice *p* = 0.71 (green p-value); interaction effect Age*Model *p* = 4.7E-8. Each data point corresponds to a linescan acquired during a single session per mouse. N = 194 APP/PS1 (55 females and 139 males), 2085 WT (947 females and 1138 males).

#### AD hallmarks and cerebrovascular dysfunction

Aβ is deposited in the brain parenchyma as neuritic plaques and along vessel walls as cerebral amyloid angiopathy (CAA), the latter exemplifying crosstalk between cerebrovascular dysfunction and AD (*24*). To probe how these pathologies relate to vascular abnormalities, we longitudinally imaged Aβ deposits labeled with Methoxy-X04 using two-photon microscopy in the same field-of-view where microvasculature was assessed (18 control and 18 AD mice, 1:1 male and female; aged 3-18 months). We quantified tissue plaque volume as well as CAA as percentage of vessel covered (Fig. 4A). Sigmoid modeling revealed similar accumulation rates (Fig. 4B), with an earlier inflection point for CAA (12.04 months) than for tissue plaques (16.75 months). Both CAA (*p* = 1.2E-12) and tissue plaque volume (*p* = 1.6E-5) were significantly correlated with decreased capillary flow rate (Fig. 4C) and lower junction density (CAA, *p* = 0.025; tissue plaques, *p* = 1.7E-5; Supplementary Fig 1).

**Figure 4.**
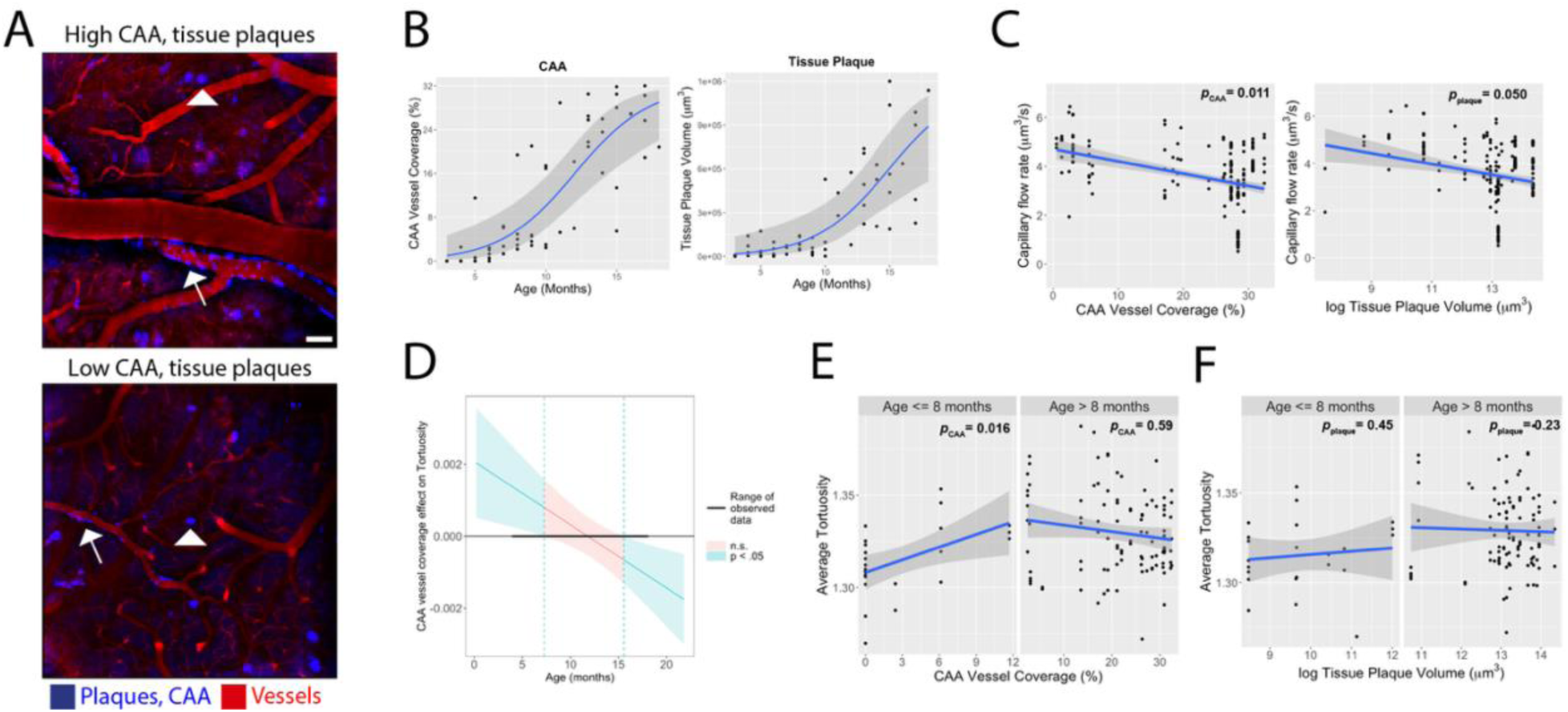
Decreased capillary blood flow rate and increased microvascular tortuosity are linked to AD pathology. A.) Amyloid-β (Aβ) deposition and vessels were labeled with Methoxy-X04 and Sulforhodamine 101, respectively (N = 9 per model and per sex, aged 3-18 months). Aβ tissue plaque volume (arrowhead) and the percent vessel coverage of cerebral amyloid angiopathy (CAA; arrow) for AD mice. Scale bar = 20 um. B.) A gaussian fit for CAA vessel coverage and tissue plaque volume versus age yielded an inflection point at 12 and 15 months, respectively. C.) Decreased capillary blood flow rate was significantly associated with both CAA vessel coverage and tissue plaque volume (linear regression analysis; CAA, *p* = 1.2E-12; tissue plaque volume, *p* = 1.6E-5). N = 223 (48 females and 175 males). D.) A Johnson-Neyman test of the interaction between CAA vessel coverage and age showed a significant association between CAA vessel coverage and microvascular tortuosity in AD mice younger than 8 months (linear regression analysis; age*CAA *p* = 0.0043) E.) Microvascular tortuosity was significantly associated with CAA vessel coverage only in AD mice younger than 8 months (linear regression analysis; age <=8 months, *p* = 0.016; age > 8 months *p* = 0.12). F.) Tissue plaque volume had no association with microvascular tortuosity across any age (linear regression analysis, age <=8 months, *p* = 0.60; age > 8 months *p* = 0.78).

We tested interaction effects between age and each Aβ pathology on microvascular tortuosity. We observed a significant interaction effect between age and CAA on average tortuosity (*p* = 0.0043). Johnson-Neyman analysis (*25*) revealed two distinct age intervals where the association between tortuosity and CAA reached statistical significance, indicating that age and CAA jointly affected vessel tortuosity in mice younger than 7.3 months and older than 15 months (Fig. 4D). Piecewise regression showed tortuosity increased with CAA up to 8 months (Fig. 4E, p=0.016). No significant interaction effect between age and tissue plaque volume on average tortuosity was observed (*p* = 0.11; Fig. 4F).

#### 7T MRI quantification of small vessel tortuosity in the human brain

Age-related changes in the tortuosity of large carotid arteries have been studied extensively with mixed results (*26*). In contrast, histological studies of small cerebral vessels (<0.5 mm) consistently report increased tortuosity with aging and AD (*27*). However, *in vivo* detection of such small vessel abnormalities has been limited by the spatial resolution of conventional 3T MRI and the inherent complexity of quantifying the morphology of higher-order cerebral vessels. We used ultra-high-field 7T TOF MR angiography with an isotropic resolution capable of capturing the small vessels (0.33 mm^3^) to investigate age-related changes in small vessel morphology in a cohort of 25 cognitively unimpaired older adults (20 female; mean age: 68.2 years). Vessel segmentation and quantification were performed using our previously developed iterative vessel tracking algorithm, VesselMapper (*28*) (Fig. 5 A, B). Vessel segments were categorized into large and small vessels using a subject-specific median split based on vessel diameter (large arteries: mean diameter = 1.85 ± 0.18 mm; small arteries: mean diameter = 1.39 ± 0.28 mm). We observed average tortuosity increased significantly with age in small arteries (*p* = 5.51E-05; Fig. 5D), but not in large arteries (*p* = 0.51; Fig. 5C). Nonlinear cross-species modeling of the human cohort and our control mice tortuosity data revealed a remarkably consistent aging pattern: small vessel tortuosity is stable through midlife and only increases in the late life stages (Fig. 5E). These findings demonstrate the strong translational relevance of our animal model results, suggesting that early changes in small vessel tortuosity may serve as a biomarker for compromised vascular health and a risk factor for late life AD.

**Figure 5.**
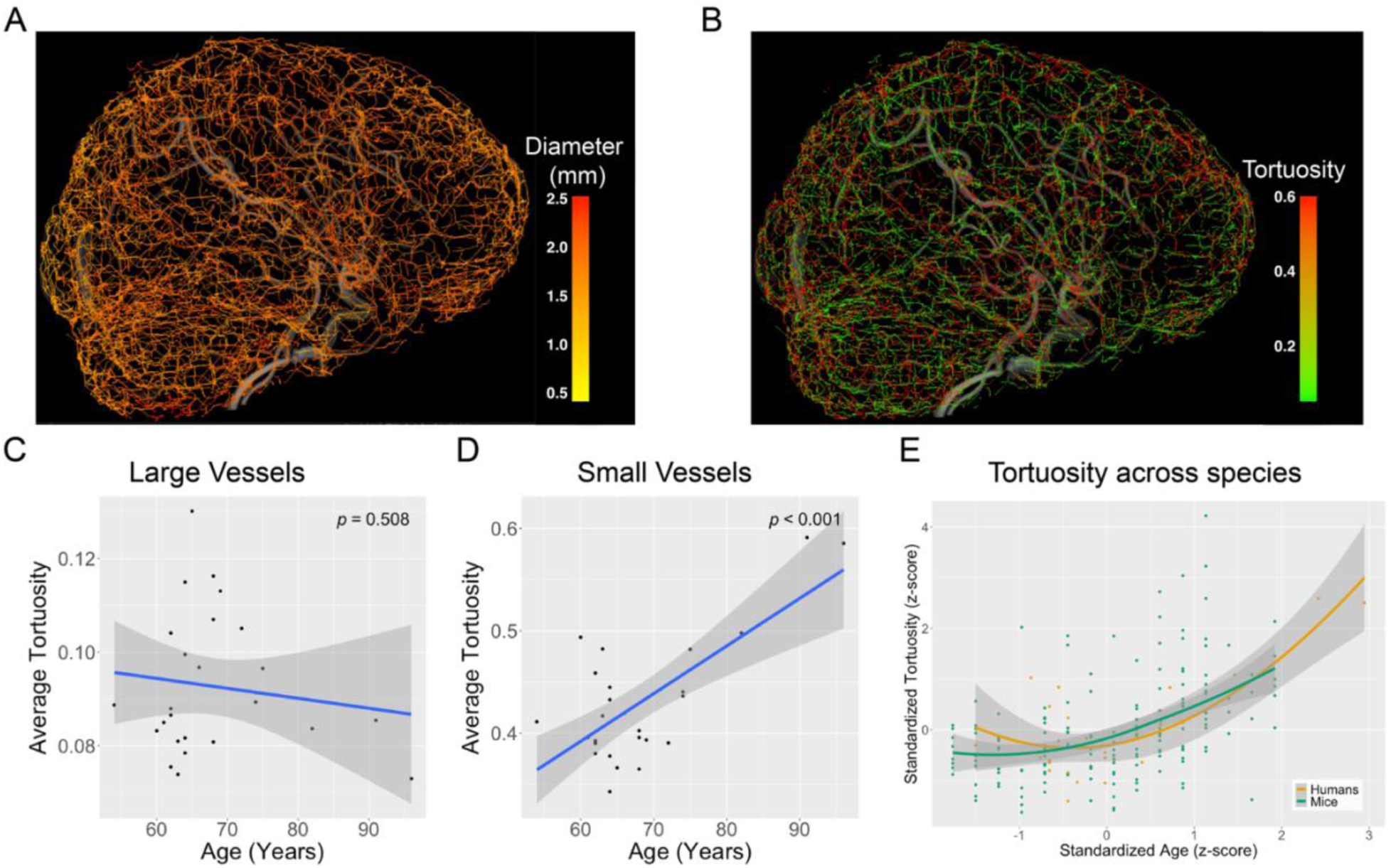
7T MRI enables high-resolution quantification of small vessel tortuosity in the human brain. Whole-brain time-of-flight (TOF) MR angiography was acquired from 25 cognitively unimpaired older adults using a 7T Siemens MRI scanner with an isotropic resolution of 0.33 mm^3^. Vessel segmentation and morphometric analysis were performed using *VesselMapper*(*28*) A.) vessel diameter and B.) vessel tortuosity across individual vessel segments. A median split based on vessel diameter was used to classify vessels as either large or small. C.) No significant association was observed between age and average tortuosity in large vessels. D.) In contrast, small vessel tortuosity increased significantly with age. E.) Z-scored LOESS-based trajectories of average small vessel tortuosity as a function of age in both control mice and the human cohort revealed a remarkably similar pattern of increased tortuosity during late life, further emphasizing cross-species translational relevance of this imaging biomarker.

#### Cerebral vessel transcriptome offers mechanistic link to vascular imaging features

To investigate the molecular mechanisms underlying our imaging findings, we performed bulk RNA-seq of isolated cerebral vessels at the 9-11 months time point marked by morphological changes (Fig. 6A). We identified 1097 differentially expressed genes between AD and controls (576 downregulated, 521 upregulated; Fig. 6B). None of the 1097 differentially expressed genes exhibited opposing expression patterns between AD males vs. control males and AD females vs. control females, suggesting shared pathways across sexes (Fig. 6C).

**Figure 6.**
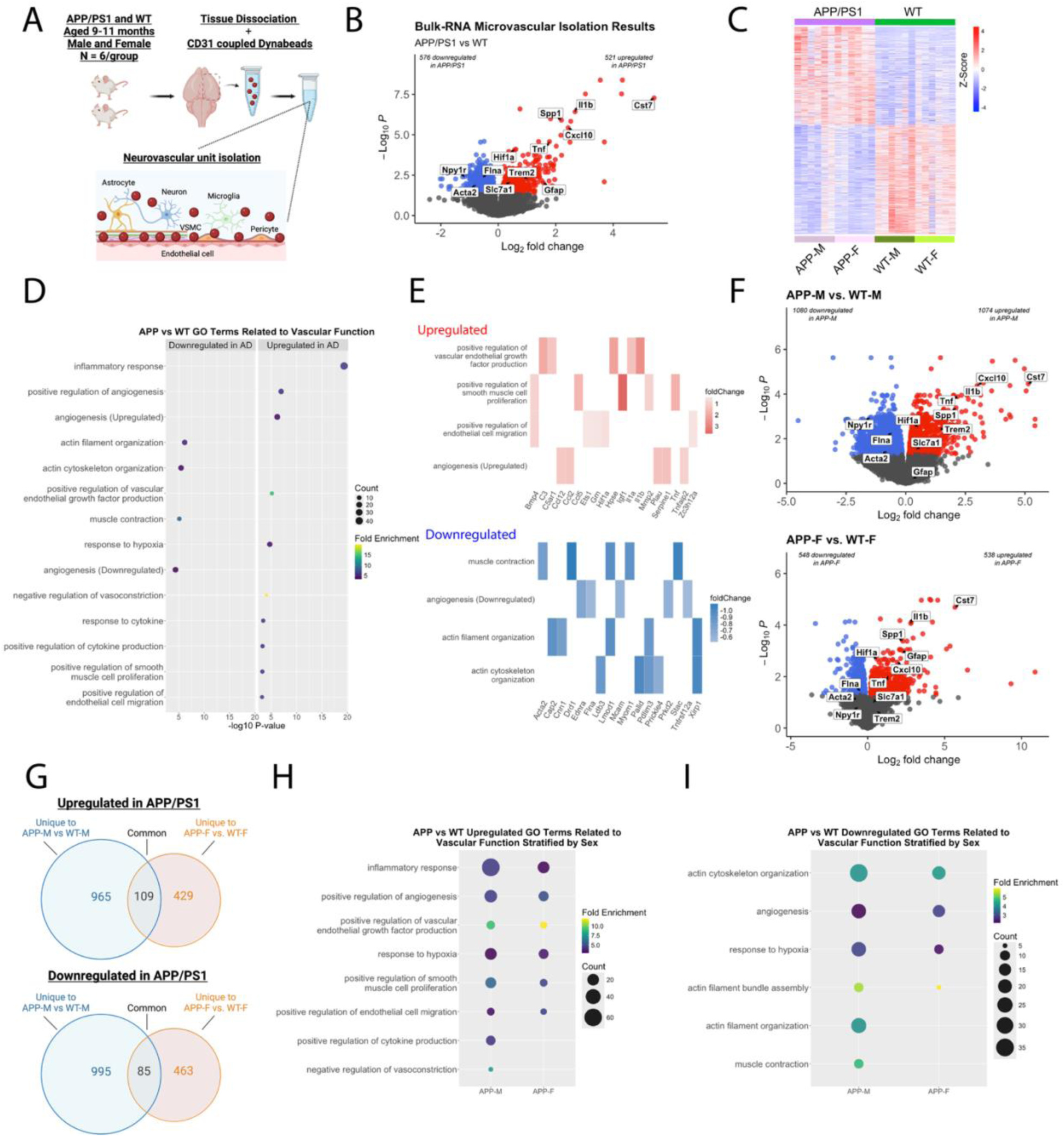
Bulk RNA-seq of isolated brain vessels in AD mice at 9-11 months of age highlight transcriptomic signatures of vascular activation and dysfunction. A.) Gene expression profiling was performed on brain vessels isolated from 9-11 month old AD and their WT controls (N = 6 per model and per sex, aged 9-11 months). B.) Volcano plot represents differentially expressed genes between AD and WT mice at FDR<0.05. Red denotes significantly upregulated genes (521 genes), blue for significantly downregulated (576 genes), and grey denotes non-significance. C.) A heatmap of DEGs at *FDR* < 0.05 displays similar gene expression patterns between sexes within a genotype for all or top-regulated genes.D.) Bubble plots showing gene ontology (GO) terms generated from either significantly down (left) or upregulated (right) genes when comparing AD and WT controls showing significant alterations in biological processes related to vascular function and structure. X axis indicates the negative log10 *p*-value, circle size is positively correlated with gene number, and circle color denotes fold enrichment. E.) The top 5 genes based on fold change which clustered within GO terms associated with vascular function are displayed for each GO term. F.) Volcano plots display DEGs for AD males vs WT males (top) and AD females vs. WT females (bottom). G.) Venn diagrams represent the common and unique DEGs between male and female mice comparisons as shown on panel F. Bubble plots display H.) upregulated and I.) downregulated GO terms for biological functions that are related to vascular function and structure, stratified by sex. X axis indicates the negative log10 *p*-value, circle size is positively correlated with gene number, and circle color denotes fold enrichment. Parts of these figures were created with Biorender.com released under a Creative Commons Attribution-NonCommercial-NoDerivs 4.0 International license.

Gene ontology analysis revealed upregulated genes clustered in neuroinflammation processes including cell migration, neutrophil chemotaxis, and interleukin-1 signaling (Fig. 6D, Supplementary Fig. 2A; Supplementary Table 1). The genes with the highest fold change were linked to neuroinflammation including *Cst7*, *Clec7a* and *Ifnb1* (*29*). Significant upregulated GO terms related to vascular function included angiogenesis, vascular endothelial growth factor (VEGF) production, response to hypoxia, smooth muscle proliferation, and endothelial cell migration (Fig. 6D). Kyoto Encyclopedia of Genes and Genomes (KEGG) pathway mapping further reflected vascular remodeling, with upregulated genes associated with fluid shear stress and atherosclerosis (Supplementary Table 1). Top upregulated genes were *Igf1* implicated in cerebral microhemorrhages (*30*) and *Spp1*/osteopontin connected to immune regulation and cell adhesion (*31*).

The 576 downregulated genes clustered in GO terms involved in actin filament/cytoskeleton organization, muscle contraction, and angiogenesis (Fig. 6D, E; Supplementary Fig. 2A, B; Supplementary Table 1). The genes with the largest fold change were *Gm3716*, *Rpl21*, and *Lao1*. KEGG pathway mapping further reflected an impaired actin-filament system, with downregulated genes associated vascular smooth muscle contraction and cytoskeleton (Supplementary Table 1). Common downregulated genes involved in actin-mediated contraction including key components of the actin filament system, such as *Myh9*, *Actn1*, *Xirp1*, and *Myh11*.

Interestingly, GO analysis revealed that both upregulated and downregulated genes clustered around processes related to angiogenesis. We observed upregulation of upstream pro-angiogenic factors associated with hypoxia and VEGF production, such as *Hif1a*, *Mmp2, Bmp4*, and *Kdr*. In contrast, we observed that downregulated genes tended to involve downstream effector pathways such as cell adhesion (e.g. *Adam15*, *Prkd2*, *Tnfrsf12a*, *Epha1*, and *Mcam)* and regulators of migration and vessel outgrowth (*32*). Together, these results build upon prior evidence of dysfunctional angiogenesis in AD (*33–35*) and coincide temporally with the emergence of microvascular tortuosity.

Given reports on sex differences in AD-related vascular dysfunction (*36*), we examined whether any of the observed processes were sex-specific. When stratified by sex, we identified 1080 downregulated genes in AD males vs. control males and 548 downregulated genes in AD females vs. control females, with 85 genes common to both comparisons (Fig. 6F, G). 1074 upregulated genes were identified in AD males vs control males and 538 in AD females vs control females, with 109 common genes. Differentially expressed genes in both sexes clustered to a number of shared GO terms indicating different mechanistic routs leading to parallel functional impairments. Upregulated genes for both sexes were enriched in processes related to immune response, angiogenesis, response to hypoxia, VEGF production, and cell migration (Fig. 6H, Supplementary Fig 2C). Downregulated genes for both sexes clustered in processes related to actin cytoskeleton organization, response to hypoxia, angiogenesis, and cell proliferation (Fig. 6I, Supplementary Fig 2C).

#### Cell-specific contributions to vascular imaging abnormalities

To identify specific cell types driving transcriptional changes, we mapped our bulk RNA-seq results to a mouse brain vascular cell-specific atlas (*17*) (Fig. 7A; Supplementary Table 3). Among the 576 downregulated genes in AD mice, 121 overlapped with vSMC marker genes, followed by 34 with vein endothelial, and 31 with pericytes. Of the 521 upregulated genes, 70 genes mapped to microglia, 43 to capillary endothelial, and 30 to brain endothelial. (Fig. 7B, Supplementary Table 3). Genes associated with vSMC showed downregulated processes related to actin-mediated cell contraction, muscle cell differentiation, and vSMC proliferation (Fig. 7C). Pericyte marker genes were enriched in downregulated processes of blood circulation, actin-mediated muscle contraction, and blood vessel diameter maintenance (Fig. 7D).

**Figure 7.**
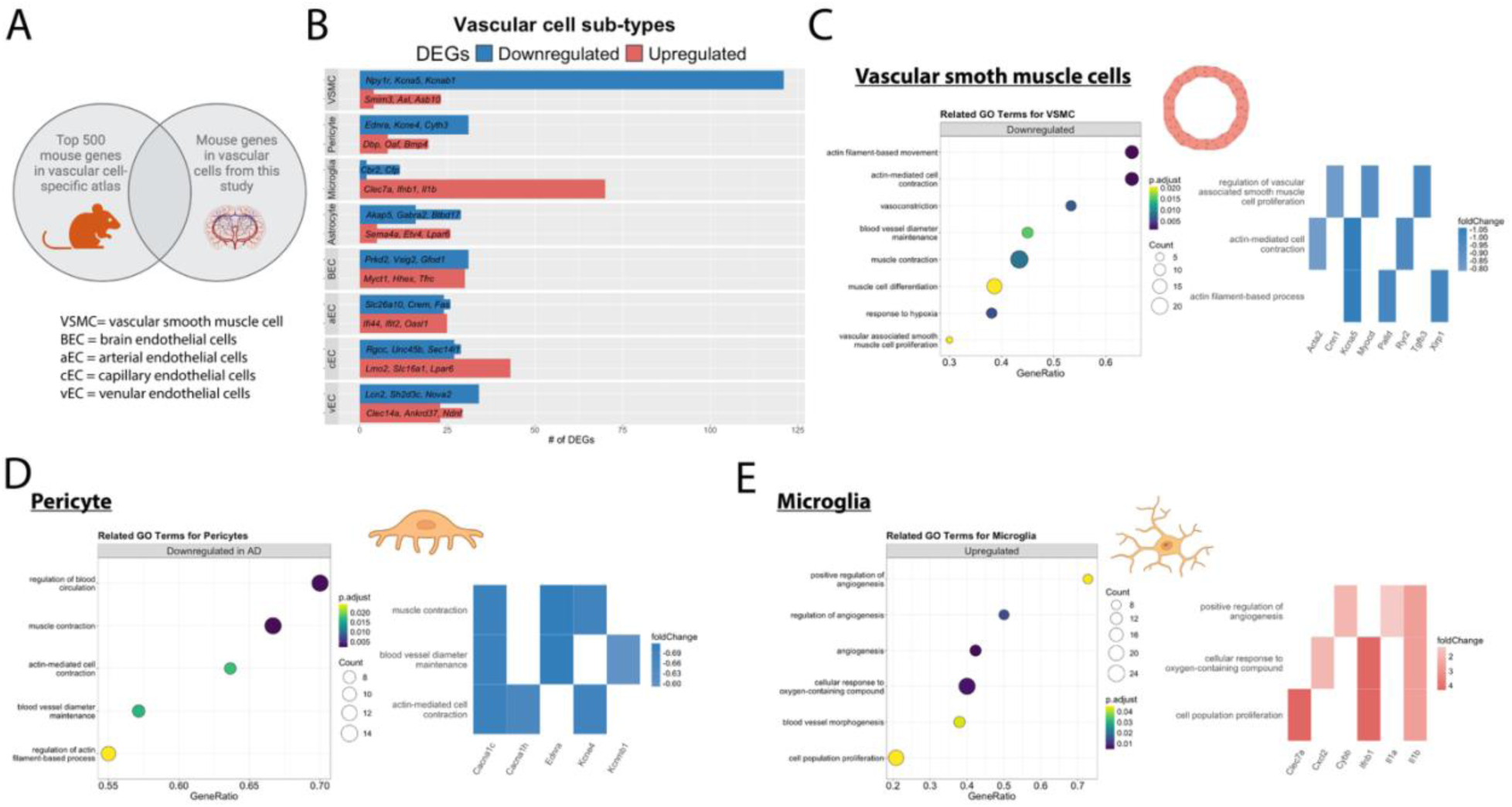
Mouse cerebrovascular cell-specific atlas implicates mural cells and microglia to impaired actin-filament system and angiogenesis, respectively. A.) Gene expression from the bulk RNA-seq on isolated brain vessels were registered to a mouse atlas(*17*) displaying the top 500 mouse brain mural cell subtype markers. B.) Bar plot showing differentially expressed genes between AD vs WT controls which show cell subtype-specific changes. C.) Bubble plot (left) showing GO enrichment analysis of vascular smooth muscle cell marker genes which display downregulated expression in vascular bulk RNAseq of AD vs WT mice are related to GO terms muscle contractility, response to hypoxia, and muscle cell proliferation. Associated heatmaps (right) displaying the top 3 genes based on fold change which clustered within vascular smooth muscle GO terms are displayed for each GO term. D.) Bubble plot (left) show GO enrichment analysis of pericyte marker genes which display downregulated expression in vascular bulk RNAseq of AD vs WT mice are related to GO terms muscle contractility and response to oxygen levels. Associated heatmaps (right) displaying the top 3 genes based on fold change which clustered within pericyte GO terms are displayed for each GO term. E.) Bubble plot (left) showing GO enrichment analysis of microglia marker genes which display upregulated expression in vascular bulk RNAseq of AD vs WT mice are related to GO terms angiogenesis and response to oxygen levels/hypoxia. Associated heatmaps (right) displaying the top 3 genes based on fold change which clustered within microglia GO terms are displayed for each GO term.

Interestingly, several genes specific to pericytes and vSMCs were associated with calcium and potassium voltage-gated channels. These ion channels are critical regulators of cerebral blood flow and implicated in AD (*37, 38*). Downregulated genes associated with calcium voltage-gated channels, *Cacna1c* and *Cacna1h*, mapped to both vSMCs and pericytes, while *Ryr2* was vSMC-specific (Fig 7C,D). Potassium voltage-gated channel genes, *Kcne4* and *Kcnbm1* mapped to pericytes and *Kcne4*, *Kcnb1*, *Kcna5*, and *Kcnab* mapped to vSMCs (Fig. 7B-D, Supplementary Table 3). These findings suggest that impaired actin-filament integrity and ion channel dysfunction in mural cells may play a significant role in the observed reduced capillary blood flow.

GO analysis of differentially expressed genes highlighted a dysfunctional angiogenic response, with upregulated pro-angiogenic factors lacking activation of downstream effectors such as cell adhesion processes (Fig. 7E). Upregulated genes also clustered around neuroinflammatory processes, implicating the involvement of microglia with markers associated with upregulated vascular-related pathways (Fig. 7E). Similarly, endothelial cells demonstrated contributions to both neuroinflammation and aberrant angiogenesis (Supplementary Fig. 3A-E). Arterial endothelial cells clustered around regulation of blood circulation and cytoskeleton organization (Supplementary Fig. 3C). Capillary endothelial cells exhibited upregulated terms related to cytokine-mediated signaling pathways and cell migration involved in sprouting angiogenesis (Supplementary Fig. 3D). Vein endothelial cells displayed upregulated terms related to inflammatory response and cell migration (Supplementary Fig. 3E). Astrocyte marker genes were enriched in processes related to ion transport and vesicle-mediated synaptic transport (Supplementary Fig. 3F).

#### Cross-species vessel transcriptome analysis

Focusing on translation, we tested whether identified mouse genes also aligned with healthy and AD human brain vascular atlas based on the same cell types (*18*). We found 18 downregulated genes from our AD mice aligned with vSMCs in humans, followed by 7 for pericytes, and 5 for astrocytes (Fig. 8A; Supplementary Table 4). Several of the vSMC and pericyte genes are linked to the actin-filament system and ion channels, including *RYR2, CACNA1C*, *KCNE4, KCNAB1, ACTA2*, *MYH11*, and *MYL9*. Among the mouse cell-specific upregulated genes, 5 genes aligned with human microglia, 4 to human capillary endothelial cell, and 2 to human vein endothelial cells. Notable capillary endothelial upregulated genes included solute carriers *SLC16A1* and *SLC7A1* which deliver lactate/pyruvate and amino acids across the blood-brain barrier (*39*).

**Figure 8.**
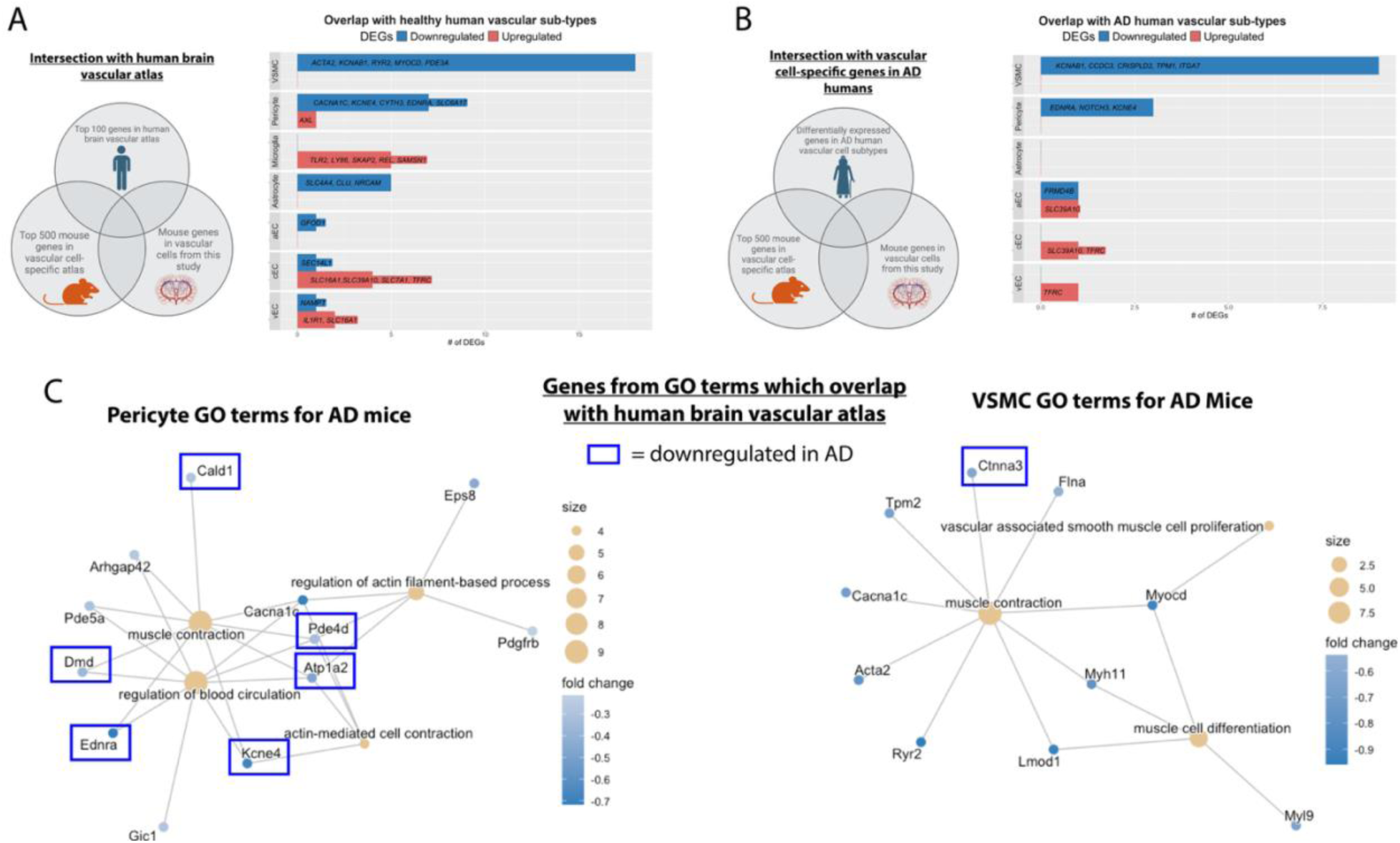
Human cerebrovascular cell-specific atlas identifies translational targets for mural cell dysfunction in AD. A.) Gene expression from the bulk RNA-seq on isolated brain vessels were mapped to a healthy human brain vascular atlas(*18*) displaying the top 100 genes in each vascular cell sub-type. Our genes were mapped to the mouse brain vascular atlas and then aligned with a healthy human brain vascular atlas(*18*). Bar plot showing number of differentially expressed genes from the bulk RNA-seq on isolated brain vessels that were also defined as in the top 100 genes in the healthy human brain vascular atlas in each vascular cell sub-type. B.) Bulk RNA microvascular isolation genes were overlapped with differentially expressed genes in AD humans for each vascular cell sub-type in the human brain vascular atlas. Our genes were mapped to the mouse brain vascular atlas and then aligned with a AD human brain vascular atlas(*18*). C.) Genes pertaining to pericyte and vascular smooth muscle cell’s GO terms which also overlapped with the healthy human vascular atlas are shown. In the blue boxes are the genes which also overlapped with differentially expressed genes for AD humans. Parts of these figures were created with Biorender.com released under a Creative Commons Attribution-NonCommercial-NoDerivs 4.0 International license.

When mapped to the AD human brain vascular atlas, 9 mouse genes overlapped with human vSMC markers, followed by 3 with pericytes, and 1 with arterial endothelial cells (Fig. 8B; Supplementary Table 4).

## Discussion

Alois Alzheimer described abnormal vessel growth and presence of vessel abnormalities in the first documented case of AD (*40, 41*). This observation was largely overlooked for decades, with recognition of early vascular abnormalities only in the past 20 years. Few have attempted to bridge how neuroimaging trajectories could be linked directly to molecular changes. Using multi-modal *in vivo* longitudinal imaging of AD mice, our study provides critical insights into cerebrovascular remodeling when pathological changes are silent. By leveraging near-lifespan imaging, we identified a critical time point when cerebrovascular abnormalities arise and used it as a road marker to isolate brain vessels for RNA sequencing. We noted inflammatory response of vessel-associated microglia, dysfunctional angiogenesis, and actin-mediated muscle contractility dysfunction. Mapping transcriptional changes to a mouse brain cell-specific atlas implicated endothelial cells and microglia in angiogenic dysfunction, while pericytes and vSMCs were linked to deficits in actin-filament system and ion channels expression. Comparisons with human brain vascular atlases identified therapeutic targets for actin-mediated muscle contractility, lactate transporters and aberrant angiogenesis. To assess whether similar early imaging features emerge in humans, we acquired 7T time-of-flight MRI in cognitively unimpaired older adults and identified significant age-related increases in small vessel tortuosity. Our human findings parallel our mouse data, supporting small vessel tortuosity as a non-invasive imaging biomarker of molecular changes associated with early AD vulnerability.

A key finding was that microvascular tortuosity increased early in AD mice. This abnormality became pronounced around 9-11 months of age and was accompanied by reduced red blood cell velocity and decreased microvascular density. At this time point, bulk RNA-seq analysis of isolated brain vessels yielded upregulation of genes involved in pro-angiogenic responses to hypoxia and VEGF signaling, alongside downregulation of downstream effector pathways, particularly cell adhesion (*32*). Vessel tortuosity is strongly linked to pathological angiogenesis at the molecular level in cancer (*42*), retinal diseases (*43*) and other brain disorders (*44*). However, AD studies traditionally focused on Aβ deposition and morphology (*45*) or transcriptomic markers of impaired angiogenesis (*33, 34*), without directly linking vascular imaging features to omics data. Our findings suggest that beyond angiogenic signaling, other pathogenic pathways likely contribute to tortuosity such as wall shear stress, endothelial and vSMC proliferation, and degeneration of the elastic layer - each likely interacting with one another (*46–48*). Given the detectability of arterial tortuosity via human MRI, further research is warranted to validate its predictive value for AD conversion in the presence of Aβ deposition and altered cerebral blood flow. Previous studies by us and others report diminished functional and hypercapnic hyperemia in AD mice with Aβ−driven arterial dysfunction (*49, 50*). In a smaller cohort of the same mouse model, we have previously shown that CAA in the arterial vessel wall is significantly correlated with red blood cell velocity (*23*). We reproduce this finding here across the lifespan, adding the significant link between CAA and vessel tortuosity. We identify downregulation of genes related to the actin-filament system, linking pericyte and vSMC dysfunction to reduced capillary blood flow. Pericytes are especially vulnerable to AD pathology, with early reductions in blood flow linked to Aβ-induced pericyte constriction (*51*) and subsequent loss of pericyte coverage (*52*). Furthermore, we identified downregulated genes associated with vascular ion channels, critical for cerebral blood flow regulation (*38*). Several of these genes mapped to human vSMCs making them attractive therapeutic targets. Indeed, voltage-gated calcium channel blockers have shown benefit in AD patients (*38*). In AD mice, blocking voltage-gated calcium channels improved blood flow (*53*). An intriguing test would be to track the vessel tortuosity alongside blocking voltage-gated calcium channels. Combined, targeting mural cell contraction early in AD may offer a viable strategy to enhance brain energy supply.

A significant finding from our sex-specific transcriptomic analysis was that minimally overlapping gene sets in male and female AD mice drove converging GO terms and similar imaging features. The important implication is that upstream vascular therapeutic targets may differ between men and women despite addressing common dysfunctions. This point is further reinforced by our recent finding in the same AD model, that while both AD males and females showed reduced vascular reactivity with age, the AD females exhibited this decrease in smaller arteries, whereas AD males experienced decreased dilation in the larger arteries (*50*). This prior finding also exposes a limitation of the present transcriptomic data coming from the full vascular tree, where small and large vessels were isolated together and classified *post factum*, thus limiting the statistical power with respect to sex differences along the vascular tree.

A major strength of our study was the integration of bulk RNA-seq data with both mouse and human brain vascular atlases, enhancing the translational relevance of our findings. However, cell-specific gene expression changes in AD may not align directly with atlases derived from healthy brain vasculature. For example, several downregulated actin-filament genes mapped to vSMCs, yet pericyte loss in AD (*52*) could be the primary driver of the observed bulk downregulation. Furthermore, the human AD vascular atlas likely reflects later disease stages, potentially explaining limited overlap with our early-stage dataset. Our findings emphasize a dynamic interplay between the inflammatory and vascular response in AD. Given that the neuroinflammatory signature of AD is stage-dependent (*54*), it is likely that the inflammatory-vascular responses also evolve throughout disease progression. While we observed promising cross-species alignment of cell contractility genes, ion channels, and lactate transporters; the mural angiogenic signatures demonstrated less overlap with human data. This discrepancy may stem from differences in disease stage or species-specific vascular cell composition. A significant translational impact of our work is quantification of the separate contributions of CAA and tissue plaques to vascular dysfunctions, because human non-invasive imaging currently can not distinguish between these two separate Aβ locales (*24*). We show that both CAA and tissue plaques contribute to decreased capillary blood flow and microvascular density, however the vessel tortuosity was highly correlated only with CAA.

Importantly, the age-related increase in small vessel tortuosity we observed in humans showed striking similarity to that in mice, reinforcing translational potential of the mechanistic routs and opening the possibility of linking proteomic markers of angiogenesis (*55*) to non-invasive imaging.

In conclusion, our study identified transcriptomic signatures underlying vascular abnormalities observed through imaging at a critical time point when vascular remodeling and dysfunction emerge early in preclinical AD model. Combined, our imaging and transcriptomic data support a model in which early CAA accumulation affects vascular remodeling and impaired angiogenic response. The associated increases in tortuosity elevate fluid shear stress that brings further remodeling, reduced vessel contractility, decreased capillary blood flow and lower tissue oxygenation.

More broadly, our results underscore the need for continued efforts to define cell-specific targets of vascular activation in the early AD in the context of MRI-based measurements of arterial tortuosity as an early biomarker. Our work integrates data from mice and humans to identify commonalities and divergences in functional and morphological vascular patterns and the underlaying cellular processes. Our objective is to capitalize on cross-species multiparametric trajectories for better diagnostics, therapy and improvement of vascular brain health of all aging humans.

## MATERIALS AND METODS

### Two-Photon Microscopy

#### Animal Preparation for two-photon imaging

All animal procedures were approved by the Division of Laboratory Animal Resources and Institutional Animal Care and Use Committee at the University of Pittsburgh and performed in accordance with the National Institutes of Health Guide for Care and Use of Laboratory Animals. We used B6.Cg-Tg(APPswe,PSEN1dE9)85Dbo mice, male and female (Jackson Labs-MMRC) transgenic for chimeric mouse/human amyloid precursor protein (Mo/HuAPP695swe) and mutant human presenilin 1 (PS1-dE9) directed to neurons (*56*). Control mice (B6C3, Jackson Labs) from the background of the AD model were age and sex matched (N = 9 AD male, 9 AD female, 9 control male, 9 control female for total of 36 mice across all groups). Prior to data collection, each animal underwent surgery at approximately 2 months of age to implant a cranial window, as previously described (*50*). Briefly, mice were anesthetized with ketamine / xylazine at 70/10 mg/kg. Body temperature was maintained at ∼37.0° C. A 4 mm craniotomy over the right primary somatosensory cortex was re-sealed with a glass coverslip, and a head-plate was fixed to the skull (Narishige, CP-2). Mice were given 4 weeks to recover and acclimated to head fixation on the treadmill, 1 hour for 5 consecutive days. All mice underwent longitudinal imaging, however not every animal contributed data at every time point, with occasional excluded sessions due to motion or attrition over time.

#### Two-Photon Microscopy Image Acquisition

One day before imaging, the mice were injected intraperitoneally with Methoxy-X04 (1 mg/kg) to visualize Aβ deposition *in vivo.* Just before imaging, the mice were injected intraperitoneally with Sulforhodamine 101 (SR101, 0.2 μl/g) to visualize the vasculature. Imaging was performed with Ultima2p microscope (Bruker Nano Inc.) coupled to tunable laser (Insight X3 dual, Newport Spectra-Physics). Awake, head-fixed mice were placed on a custom-made frame with movable horizontal drums designed to accommodate mouse locomotion. Images of the vasculature and Aβ deposition were acquired using 16x water immersion objective (0.80 NA, Nikon), with excitation at 920 nm and 740 nm. Z-stack volumes over somatosensory and motor cortex were obtained with a field-of-view of 1,130 x 1,130 μm (512 × 512 pixels), at up to 400 μm cortical depth. This corresponds to an in-plane resolution of 2.2 μm/pixel with a step size of 3 μm between xy image planes. Line scans (acquisition speed 833.34 Hz; pixel size 0.37 μm) were also taken from individual blood vessels to quantify vessel diameter and red blood cell velocity.

### Two-Photon Microscopy Image Analysis

#### Vessel Morphology Segmentation and Quantification

First, vascular trees were semi-automatically segmented using ilastik (*57*). Utilizing the MONAI framework (*58*), we trained a 5-layer residual U-net segmentation model (*59*). Total of 15 images were used for training and 5 for validation. The model was trained with a learning rate of 1e-4, batch size of 2, and the Adam optimizer. To augment the training dataset, we applied random rotations, flipping, and volumetric cropping to dimensions of 256 × 256 × 32 pixels. The network was trained for 600 epochs using the Dice loss function, achieving a final Dice score of 0.929. Once segmented, images were skeletonized via medial axis thinning. The skeletonized vascular trees were imported into Fiji ImageJ (*60*) skeleton analysis yielding measurements of individual endpoints, junctions, and branch lengths (*61*). The tortuosity index was calculated as the skeleton arc length divided by the corresponding Euclidean distance between junctions or junction-endpoints. The density of the vasculature was measured by normalizing total junctions by the volume in the top 50 slices where the intensity distribution was constant. Vessel segmentations were visualized with VesselVio (*62*).

#### Red Blood Cell Velocity Analysis

All data were analyzed using MATLAB (MathWorks) as previously described (*23*). Briefly, line-scan data was obtained by repetitive scanning along a center-line segment of 2-10 capillaries per imaging session. The raw data strips were pre-processed into blocks of maximum length 250ms. We applied Radon transform to estimate the angle of maximum variance for velocity estimation, which was further refined based on signal-to-noise ratio (SNR), yielding the estimated velocity for each data block (*22*). The vessel diameter was determined by calculating the full-width-at-half maximum of a rectangular vessel cross-section.

#### Cerebral amyloid angiopathy (CAA) and plaque volume quantification

All data were analyzed using MATLAB (MathWorks, MA, USA). The two-photon z-stacks were first pre-processed by performing median filtering (3-by-3 neighborhood), gaussian smoothing (σ = 0.4), and removing the low-frequency intensity fluctuations extracted from the average image. Volume intensity correction was performed in the z-direction where each quantile of the volume was normalized to the signal intensity of the surface quantile. For CAA quantification, vessel wall Aβ deposition was identified from the overlap with the SR101 vascular channel on a maximum intensity projection (MIP). Vessels with CAA were manually segmented from the MIP, and pixels categorized as CAA were determined by thresholding 1.5 standard deviations above the mean intensity of Methoxy X04 in vessel, then the percentage vessel coverage of CAA with respect to the total vessel area was computed. Aβ tissue plaque quantification started with the removal of CAA pixels from the volume followed by segmentation of plaques based on a threshold of greater than 0.6 times the maximum intensity. Tissue plaques were identified as segmented pixels with connected components (8-connected) greater than 15 pixels, and the total volume was calculated from the resulting pixels. Measures of CAA vessel coverage and tissue plaque volume were averaged over the imaging session to compute a global average for each time point in each mouse.

#### Mouse MRI Acquisition and Image Analysis

##### Animal Preparation

Independent from the two-photon imaging cohort with aluminum chronic optical windows, we performed MRI scans on isogenic AD mice (N = 47, 23 male and 24 female) and the B6C3 control mice (N = 17, 8 male and 9 female; aged 2-25 months). Mice were anesthetized with isoflurane through scanning. The dynamic respiration rate and end-tidal CO2 were monitored (Biopac). To ensure stable animal physiology during the MRI experiments, isoflurane was adjusted in the range of 1% to 1.5% to keep the respiration rate below 90 breaths per minute. The body temperature was kept at 37.5±1 °C using a rectal probe feedback-controlled water-exchange heating bed integrated with the MRI cradle.

##### MRI Acquisition

All MRI scans are performed on a 9.4T horizonal bore with 86mm volume coil for transmit and a 4 channel mouse head RF coil for receive (Bruker). Mice underwent MR-imaging sessions to acquire TOF angiographies and T2-weighted anatomical reference scans. 3D TOF MRI was acquired with 3D gradient echo Fast Low Angle Shot (FLASH) with (echo time, TE= 2 ms and repetition time, TR= 12 ms). The full brain field of view (FOV) was 20 x 20 x 15 mm with a matrix size of 256 x 256 x 192 voxels. This resulted in isotropic voxel size of 78 microns across the whole brain.

##### TOF MRI Image Analysis

TOF images were first skull stripped with FSL Brain Extraction Tool and manual touch-ups. The MRI images were segmented using exactly the same semi-automatic approach described above for two-photon vessel segmentation. We also used exactly the same approach as described above to skeletonize the segmentation, quantify vessel tortuosity and junction density and visualize the resulting full brain vascular trees. Using the T2-weighted MRI, images were co-registered to a B6C3 mouse MRI atlas with AFNI (*20, 21*).

#### Human MRI Acquisition and Image Analysis

##### MRI Data Acquisition

Data was collected on 25 cognitively normal older adults (20 female, mean age 68.2 years) from an ongoing study described previously (*63*). MR data were acquired on a 7T Siemens Magnetom scanner on a custom 16-transmit and 32-receive channels (Tic-Tac-Toe design) RF coil (*64*) operating in the single transmit mode. Structural images were acquired using a 3D multiecho Magnetization Prepared Rapid Gradient Echo (MPRAGE) sequence (TR = 3000ms, TE = 2.17 ms, TI = 1200 ms, 0.75 mm^3^ isotropic resolution, and acceleration factor of 2). TOF images covering the whole brain were acquired using a 7-slab protocol covering the whole brain (TR = 14 ms, TE = 4.5 ms, resolution = 0.38 mm^3^, 354 slices, and slab overlap = 17.86 mm).

##### TOF MRI Analysis

Vessel segmentation and morphometric analysis were performed using *VesselMapper*, an automated pipeline implemented in Python 3 and ITK 5.3 (www.itk.org), as described previously (*28*). Briefly, vesselness probability was computed for each voxel using the Hessian matrix, enabling the identification of tubular structures representing arterial vessels. Connected components were extracted from the vessel maps. To address discontinuities and local artifacts, an iterative skeletonization and refinement approach was employed to estimate the vessel medial axis and identify junctions and endpoints. Vessel paths between the junctions were reconstructed using a restricted flood fill algorithm. The final output produced an accurate skeletonized map of the vascular network. From this skeletonized representation, key morphometric features such as vessel diameter and tortuosity were extracted.

#### Statistical Analysis for Imaging Data

All statistical analyses were performed using R (version 4.3.1 https://www.R-project.org). Linear regression models were used to assess the effects of age, sex, genotype (AD vs. control), and their interactions on measured outcomes across all experiments. For group comparisons, ANOVA analyses followed by Tukey’s multiple comparison test were performed. The Johnson-Neyman test was utilized to examine the conditional effect of age on average tortuosity versus CAA for AD mice (*25*). All researchers were blinded to experimental groups during the analysis. All results are reported as means ± SEM. Number of experiments and statistical information are stated in the corresponding figure legends. In figures, asterisks denote statistical significance marked by *p < 0.05; **p < 0.01; ***p < 0.001.

#### Bulk RNA Sequencing of Isolated Mouse Brain Vasculature

##### Brain Vessel RNA isolation

Brain vessel isolation was performed as previously documented with slight modifications (*65*). Mice (9-11 months of age, 6 mice per sex and phenotype, total of 24 mice) were euthanized and brain removed from the skull. The brain was placed in ice cold primary washing medium with 1% Penicillin/Streptomycin and cut into 1 mm^3^ pieces. Tissue pieces were dissociated in 3 ml freshly prepared dissociation buffer (Sigma). The solution was transferred to a new tube and placed in a rotating incubator (speed 20 rev/min) at 37°C for 10 min. Samples were filtered through a 70mm mesh strainer (Falcon) to remove large fragments. The filter was washed with 10ml of primary washing medium, and spun at 600g for 5 min. The supernatant was removed, and pellet was divided into three tubes. Rat anti-mouse CD31 antibody (BD, 10 µl/brain) coupled to Dynabeads® Sheep Anti-Rat IgG (Invitrogen) were added to each sample and incubated for 1 hour at room temperature. The supernatant was replaced with 1.5ml freshly made secondary dissociation buffer (10x TrypLE with 1 mg/ml collagenase IV, Gibco). Samples were placed in rotating incubator for 5 min at 37°C, slowly resuspended ten times with a 20-gauge needle and then beads were captured on a magnetic stand. The supernatant containing the dissociated cells was washed with ice cold PBS and centrifuged for 5 min at 3,000 g (4°C).

##### RNA Isolation and sequencing

Total RNA was extracted using the RNeasy Mini kit (Qiagen) and integrity measured with an Agilent RNA Pico Bioanalyzer assay (Agilent Technologies). Libraries were generated using TruSeq standard mRNA kit (Illumina) according to manufacture’s instruction. Each sample received a single index/barcode during the library generation. Finally, the library quality was validated with a High Sensitivity DNA chip on Agilent 21000 Bioanalyzer. Sequencing was performed at the Health Sciences Sequencing Core at UPMC Children’s Hospital of Pittsburgh on NextSeq 2000 (Illumina).

##### RNA-seq Data Analysis

The sequencing data was aligned to the mouse genome using Rsubread (v2.16.1) with an average read depth of 150 million successfully aligned reads. To calculate fold changes and FDR-corrected p-values, statistical analysis was carried out using Rsubread (v2.18.0), DEseq2 (1.44.0), and EdgeR (v4.2.1), in R environment (v4.4.1). To identify biological processes associated with significantly affected genes in the AD compared to control mice, Gene Ontology (GO) and KEGG pathway analysis was performed with Database for Annotation, Visualization and Integrated Discovery (DAVID, version 6.7)(*66*). Genes identified from isolated brain vessels were compared to mouse atlas displaying the top 500 mouse brain vascular cell markers (*17*). Gene Set Enrichment Analysis of GO biological processes used the Bioconductor package clusterProfiler (version 4.12.6). Because of the substantial reduction in the number of overall genes once mapped to a mouse atlas, we analyzed GO terms from gene set enrichment analysis for each cell-type that had an uncorrected *p*-value < 0.05. Genes which mapped to the mouse brain vascular atlas were overlapped with the top 100 genes in each vascular cell sub-type in a healthy human brain vascular atlas and corresponding differentially expressed genes in AD humans (*18*).

## Data and resource availability

The datasets generated during and/or analyzed in the current study are available from the corresponding author upon reasonable request. Code for processing imaging data in figures 1 through 4 can be found at https://github.com/TILabPitt/VesselMorphologyTrajectories.

## Funding and assistance

This work was supported by the National Institute of Health, grant number RF1NS116450 and R01AG092661 (B.I), T32MH119168 (N.S.), R01AG085566 and R01MH122604 (MW), R01AG075069 (N.F.F), and R01AG077636 and RF1AG075992 (R.K.).

## Author contributions and guarantor statement

B.I. and N.F.F are the guarantors of this work and, as such, had full access to all the data in the study and takes responsibility for the integrity of the data and the accuracy of the data analysis.

