## Supplementary Figures for "Linking cross-species trajectories of cerebrovascular remodeling in aging and Alzheimer’s disease to brain vessel transcriptome"

Noah Schweitzer *et al*

**This PDF file includes:**

Supplementary Figures
Figs. S1 to S3

**Other Supplementary Materials for this manuscript include the following:**
Data S1 to S4

### Supplementary Figures

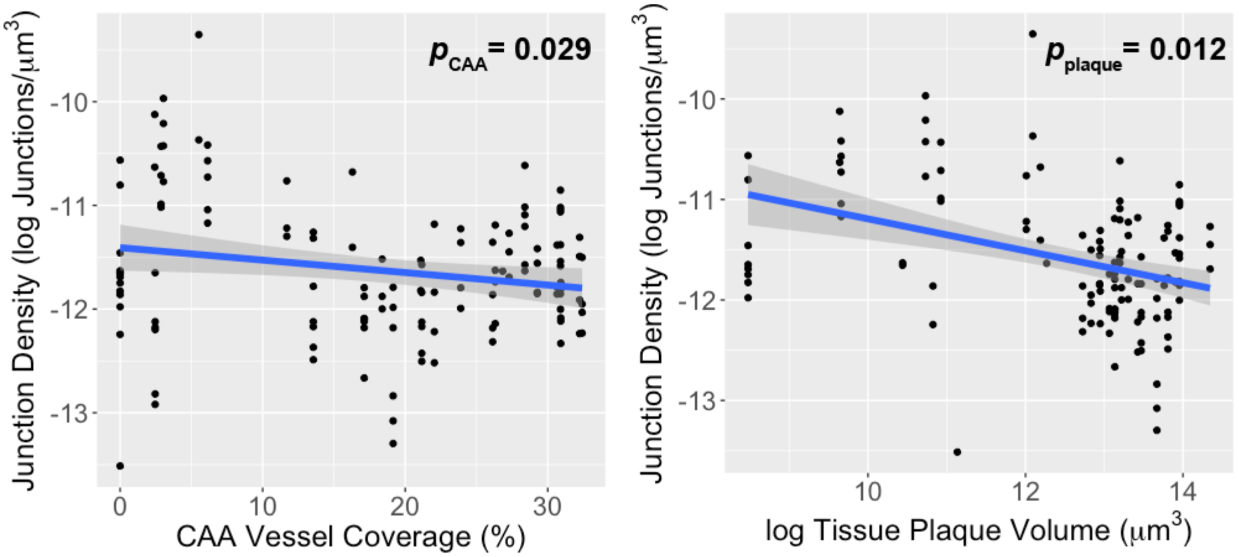


**Supplementary Figure 1. Microvascular junction density is significantly associated with both CAA vessel coverage and tissue plaque volume.** Each data point corresponds to a 3D two-photon image volume acquired during a single session per mouse. Linear mixed effect model with mouse as a random effect was performed to calculate p values.

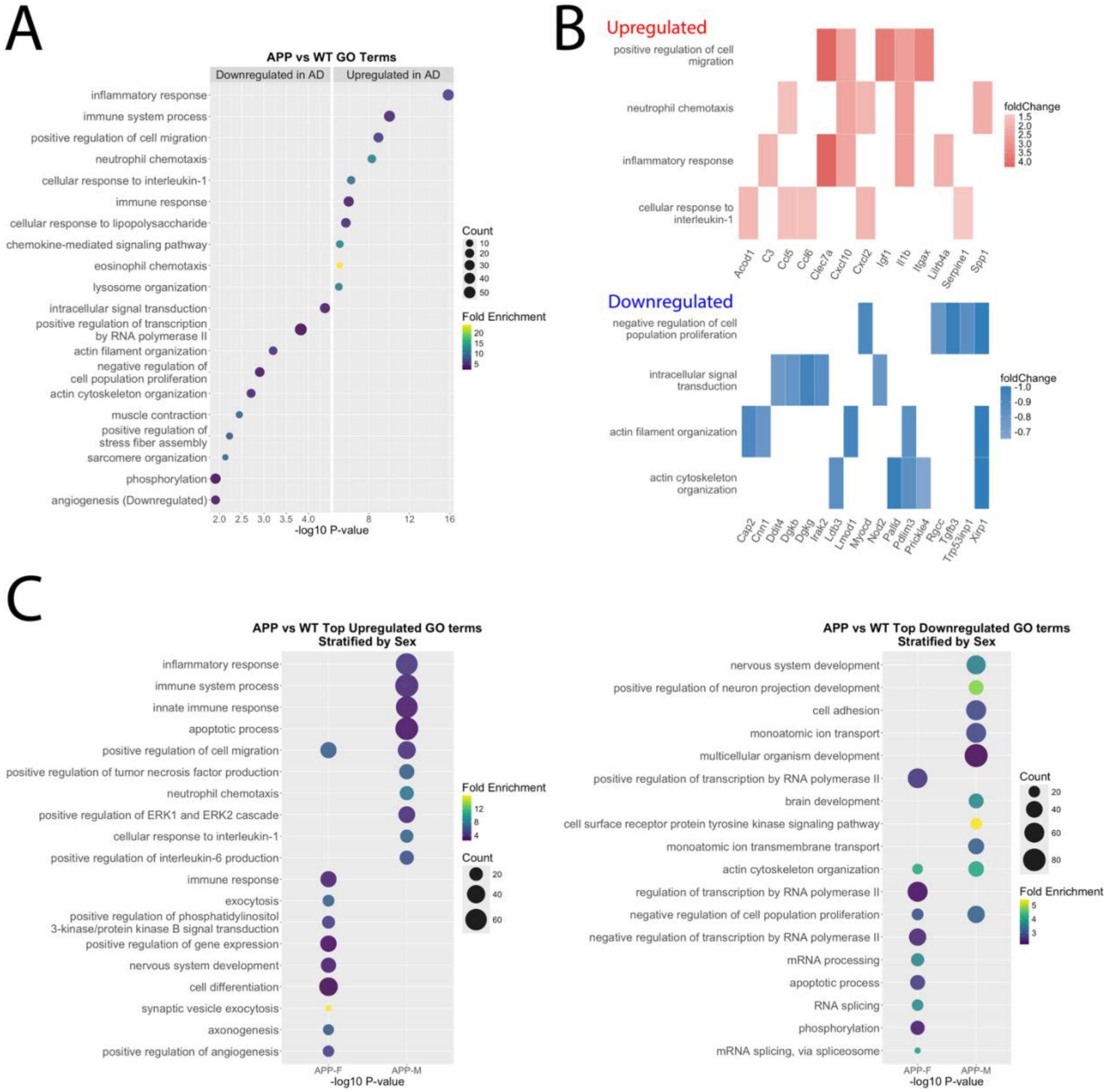


**Supplementary Figure 2. Top GO terms from bulk RNA brain vessel isolation implicate heightened immune response and diminished blood flow regulation.** A.) The GO terms with the 10 lowest p-values for all AD mice are displayed. B.) The top 5 genes based on fold change which clustered within top GO terms are displayed for each GO term. C.) The bubble plot denotes GO terms with the 10 lowest p-value in each genotype from both upregulated and downregulated DEGs. X axis indicates the negative log10 *p*-value, circle size is positively correlated with gene number, and circle color denotes fold enrichment.

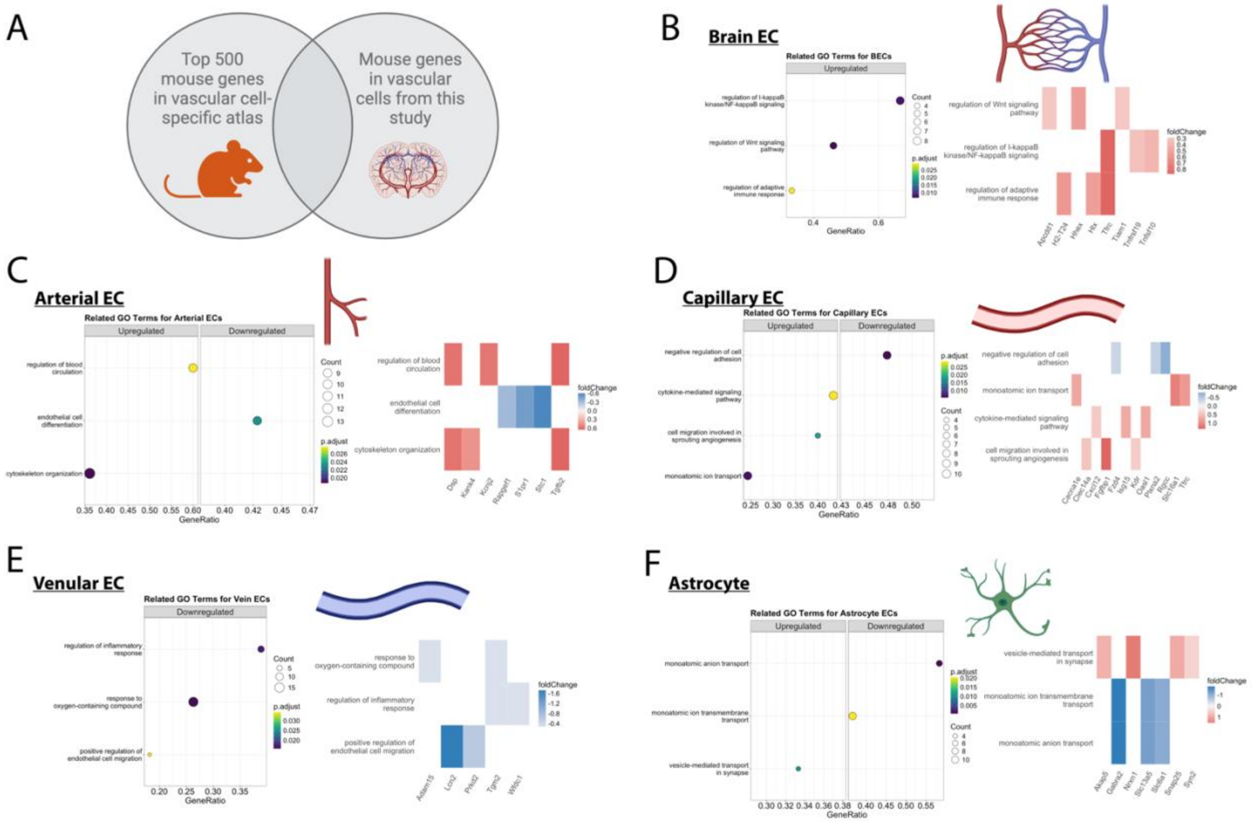

**Supplementary Figure 3. Mouse cerebrovascular cell-specific atlas implicates endothelial cells to processes related angiogenesis and proliferation.** A.) Gene expression from the bulk RNA-seq on isolated brain vessels were mapped to a mouse atlas[31] displaying the top 500 mouse brain mural cell subtype markers. B.) Bubble plot (left) showing GO enrichment analysis of all brain endothelial cell marker genes which display upregulated expression in vascular bulk RNAseq of APP vs WT mice are related to GO terms cell proliferation and blood brain barrier transport. Associated heatmaps (right) displaying the top 3 genes based on fold change which clustered within brain endothelial cell GO terms are displayed for each GO term. C.) Bubble plot (left) showing GO enrichment analysis of arterial endothelial cell genes which display upregulated expression in vascular bulk RNAseq of APP vs WT mice are related to GO terms cell proliferation and regulation of angiogenesis. Associated heatmaps (right) displaying the top 3 genes based on fold change which clustered within arterial endothelial cell GO terms are displayed for each GO term. D.) Bubble plot (left) showing GO enrichment analysis of all capillary endothelial cell genes which display expression in vascular bulk RNAseq of APP vs WT mice are related to upregulated GO terms angiogenesis and downregulated GO terms related to regulation of phosphorous metabolic processes. Associated heatmaps (right) displaying the top 3 genes based on fold change which clustered within capillary endothelial cell GO terms are displayed for each GO term. E.) Bubble plot (left) showing GO enrichment analysis of venular endothelial cell genes which display expression in vascular bulk RNAseq of APP vs WT mice are upregulated GO terms related to blood brain barrier transport and downregulated GO terms related to actin-filament processes. Associated heatmaps (right) displaying the top 3 genes based on fold change which clustered within venular endothelial cell GO terms are displayed for each GO term. F.) Bubble plot (left) showing GO enrichment analysis of astrocyte cell genes which display upregulated expression in vascular bulk RNAseq of APP vs WT mice are related to GO terms monoatomic anion transport. For all bubble plots, X axis indicates the negative log10 *p*-value, circle size is positively correlated with gene number, and circle color denotes fold enrichment. Parts of these figures were created with Biorender.com released under a Creative Commons Attribution-NonCommercial-NoDerivs 4.0 International license.
